# Dispersal mediates metapopulation response to local and regional stressors

**DOI:** 10.1101/2025.09.17.676929

**Authors:** Christopher A. Halsch, Matthew L. Forister, Eliza M. Grames

## Abstract

Many populations face multiple Anthropogenic threats simultaneously, and as a result, we have observed loss of biodiversity worldwide. Moreover, population stressors act at different spatial scales, and the understanding of variation in outcomes for species will depend on their dispersal ability. In this study, we developed a spatially explicit metapopulation simulation to investigate how stressors that act at different spatial scales interact with landscape composition and dispersal behavior to drive patterns of metapopulation extirpations. We are particularly interested in gaining insight into the decline of not only range-limited species, but also widespread butterflies that have been reported in recent years, contrary to conventional wisdom about traits that make species more at risk of population decline. We found that stressors acting at different spatial scales interact with dispersal, especially in highly developed landscapes. On average, being less dispersive produces worse outcomes because more dispersive species benefit from semi-natural habitats, which can strengthen connections between source populations. At the same time, it is the degradation of these types of land in particular that may disproportionately impact dispersive species. These findings enhance our understanding of insect biodiversity loss and demonstrate that the conservation of widespread insects will likely require consideration of larger-scale landscape connectivity.

## Introduction

From an ecological perspective, the past century can only be characterized as a period of unprecedented change (Dirzo et al. 2014, Wagner et al. 2021, Edwards et al. 2025). Large-scale modification and destruction of natural lands have significantly reduced global habitat availability, and remain one of the primary drivers of the global biodiversity crisis (Raven and Wagner 2021, Caro et al. 2022, Halsch et al. 2025). Landscape-level stressors are often accompanied by, or are the cause of, additional local stressors, such as agrochemical pollutants or the introduction of non-native species, which can have further and synergistic negative impacts on population vital rates. As the universal threat of climate change continues to escalate, the potential for combinatory effects among Anthropogenic stressors is almost limitless (Parmesan and Yohe 2003, Staudt et al. 2013, Harvey et al. 2023). Populations facing multiple stressors in increasingly fragmented landscapes are likely to decline, but how stressors that act at different spatial and temporal scales interact to drive declines is uncertain and difficult to measure in the field, a problem that is exacerbated by the lack of a clear theoretical framework for understanding interacting stressors across scales (Brown et al. 2013).

Recent years have seen rapid growth in the attention paid to invertebrate population trends (Althaus et al. 2021, Wagner et al. 2021). Species like the monarch butterfly (*Danaus plexippus*) and Franklin’s bumblebee (*Bombus franklini*) have galvanized the public and become symbols of conservation and land stewardship (Thorp 2005, Preston et al. 2021). The monarch in particular demonstrates a relatively under-appreciated feature of insect declines: many declining species are widely distributed (Van Dyck et al. 2009, Wagner 2020). For example, the west coast lady (*Vanessa annabella*), which has a distribution that covers the entire western United States, is the butterfly in steepest decline across that region (Forister et al. 2021, 2023b). Traditionally, insect conservation has focused on range-limited species, often characterized by low dispersal capabilities, with only a few populations remaining (Murphy and Weiss 1988, Wagner and Van Driesche 2010, Marschalek and Klein 2010). While protecting these remnant populations will continue to be of paramount importance for conservation, the decline of widespread species is also of great concern, particularly because the spatial scale at which conservation actions need to be taken is likely very different from that of geographically limited species.

Widespread insects differ from geographically limited species in important ways. They are more likely to be dietary or habitat generalists and can occupy a greater proportion of the landscape, including degraded or suboptimal areas (Bender et al. 1998, Devictor et al. 2008). Widespread species are also more influenced by the greater landscape context, and presence and abundance at any one site can depend on processes occurring at that site, as well as on ecological interactions or environmental conditions occurring far away. In some extreme cases, the number of migratory insects in one location can vary from year to year by orders of magnitude, depending on overwintering conditions thousands of miles away (Hu et al. 2021). Large-scale spatial dependency is also true for widespread non-migratory insects that spend all year in a landscape and have high dispersal ability (Pardikes et al. 2017). Thus, as habitats continue to be degraded or lost, widespread species may be especially affected, and we might expect that species with broader geographic distributions are just as likely to decline as those with more isolated distributions.

Insight into the effects of habitat loss and degradation on widespread species can be found in the meta-population literature (Vance 1984, Vogwill et al. 2009, Wang et al. 2015). Increased dispersal can stabilize population networks, reduce population variability, and facilitate population rescue through demographic input and recolonization of extirpated areas (Lande et al. 1998, Johst et al. 2002). Yet, outcomes of high dispersal are not universally positive. Greater dispersal can lead to increased synchrony among populations, making the meta-population more susceptible to large-scale disturbance (Heino et al. 1997, Dey and Joshi 2006, Haynes and Walter 2022). High emigration rates can also have adverse effects in areas with small populations or small habitat patches due to the direct loss of reproducing individuals (Hanski 1998).

Approaches to studying these questions have often been theoretical, although one meta-analysis using butterfly data has shown a negative relationship between population growth and dispersal (Baguette and Schtickzelle 2006). The relationship between dispersal, landscape configuration, and population persistence is likely highly context-dependent and varies between landscapes, stressors, and the inherent biology of the study organisms (Johst et al. 2002).

In this study, we employ a spatially explicit metapopulation simulation to investigate the effects of multiple population stressors operating at different spatial scales, as well as variation in dispersal behavior, on metapopulation dynamics. Our goal is to understand how varying intensities of local and landscape-wide stressors interact to drive population equilibria in a landscape with different distributions of habitat quality. We are motivated in particular by the phenomenon of declining widespread butterflies from the California Central Valley (Shapiro, Arthur 2024), a highly converted landscape (Forister et al. 2010, Halsch et al. 2020). First, we ask how metapopulations respond to landscape effects when synthetic species have different dispersal capabilities. Next, we consider whether additional stressors, acting at local and regional scales, have a greater impact on metapopulation persistence and examine how stressors interact with dispersal distance. Finally, we ask how the composition of simulated landscape conservation interacts with simulated stressors to aid in the recovery of dispersive species.

## Methods

### Overview

The metapopulation simulation consisted of three stages run for 70 time steps (20-year burn-in followed by a 50-year observation period). First, a 100 x 100 cell landscape was randomly generated with varying proportions of natural, semi-natural, and developed habitats, as well as different levels of patchiness. Within each time step of the simulation, the population growth rate in each cell was influenced by environmental factors, including habitat quality, landscape-wide stress, local stress, and density dependence. Individuals then dispersed, with both immigration to and emigration from a patch coinciding. At the end of each simulation iteration, the average abundance and extinction dynamics were summarized. Parameters were varied for the magnitude of landscape-wide stress, local stress, and dispersal ability, and simulations were run for all combinations of these environmental stressors and dispersal parameters. Inference is not drawn from temporal trends within a single iteration of the simulation, but rather from comparing summary statistics among iterations with different parameters. All simulations were run using R version 4.5.1 (R Core Team 2023).

### Annual population growth rate

The primary variable affected across different conditions of the simulation was the annual population growth rate (Sibly et al. 2002). In the initial time step, populations were given a size proportional to the habitat quality in each cell. Subsequently, the growth rate for a time step was the sum of the impact of landscape, weather, additional stressors (local and landscape-wide), and density dependence based on abundance in the previous year. These effects were summed to determine the population growth for that time step (positive effects had positive values and negative effects had negative values). This sum was then exponentiated and multiplied by the previous year’s population size. Finally, this value was used as the rate parameter in a single draw from a Poisson distribution, whose value became the population size in the current step (before dispersal). The density dependence function was a right-skewed function, where middle values are assigned positive population growth, while high and very low values (Allee effects) are assigned negative population growth (Fig. S1). All other effects are described in detail in subsequent sections.

### Generating baseline habitat quality and patchiness

Landscapes were created by first generating an empty 100 x 100 matrix. A determined number (based on an input parameter) of randomly selected cells were classified into one of three land types: natural, semi-natural, or developed. The more cells that were initially placed, the patchier the resulting landscape. The landscape was then grown from these initial patches, with neighboring cells assigned to the same class as their nearest neighbor until it reached other cells that had already been assigned to a land use type. The final cells that bordered two different land use types were assigned probabilistically based on the composition of the surrounding landscape. We also determined the initial proportion of cells assigned to each land use type as another parameter of interest. The resulting landscape after this process was a matrix composed of three land-use types, varying in patchiness and proportions based on the starting parameters. This landscape generation process is random, and it can result in landscapes that do not reflect the desired land cover proportions. Because of this, we iteratively generated landscapes and included only those in the simulation where the final percent cover of land use types was within 1% of the target percent cover initially specified for each land use type. We varied both the patchiness and the percent cover of each land cover type as documented in Table 1.

**Table 1.** Description of all varied parameters in the simulation. Multiple iterations were run for all parameter combinations. The number of iterations for each parameter combination was determined from a power analysis (described in the supplement).

| Parameter | Values | Description |
| --- | --- | --- |
| Landscape composition | 5%, 16%, 25%, 33%, 50%, 66%, 90% cover | The percent cover of the landscape of each habitat type. |
| Landscape patchiness | 10, 200, 400 patches | The number of initial patches in landscape creation (shown in Fig. 1). |
| Local effect | -1, -0.75, -0.5, -0.25, 0 | The additional local effects associated with patches in semi-natural habitats. |
| Regional effect | 0, 0.25, 0.5, 0.75, 1 | The magnitude of the worsening of regional weather effects. |
| Dispersal distance | 1, 3, 5, 8, 10 cells | The number of cells away that individuals may move to during one time step. |
| Landscape restoration | 10%, 20%, 30% modified | The percent of the landscape that is changed from developed to natural. These values were used for both types restoration simulations. |

Next, each land use type was assigned a different baseline habitat quality. The quality of each natural habitat cell was drawn from a normal distribution with a mean of 0.5 and a standard deviation of 0.1. The quality of each semi-natural cell was drawn from a normal distribution with a mean of −0.25 and a standard deviation of 0.1. These values are later exponentiated as part of the process that determines annual population growth. Done this way, natural habitats were consistently a source, while semi-natural habitats, on average, were sinks, but could switch to sources if conditions were otherwise favorable. Cells classified as developed were assigned a value of −5, ensuring that even the best combinations of all other parameters could not render it viable. The most important aspect of the assigned habitat values was not the value itself, but rather how it compared to the other effects in the simulation. By choosing evenly divisible values and keeping all parameters in a comparable range, we examined how land cover compared with other additional stressors.

Finally, each landscape was given edge effects, where cells were adjusted based on the surrounding landscape composition. This was done by taking the mean habitat quality value of the 3×3 neighborhood around a cell, multiplying that value by 0.5 (to reduce its importance relative to the primary cell), and adding that value to the habitat quality value of the cell. This resulted in smoothed landscapes with varying patchiness, habitat composition, and additional local stressors. The simulation was run on ten randomly generated landscapes for most parameter combinations. However, more landscapes were used when the proportion of semi-natural habitat was higher, as a preliminary analysis indicated greater uncertainty in these landscapes (see the supplement for a complete description of the preliminary analysis, Fig. S2, S3). Examples of landscapes are in Figure 1 and an online tool described in the supplementary materials (Fig. 1).

**Figure 1.**
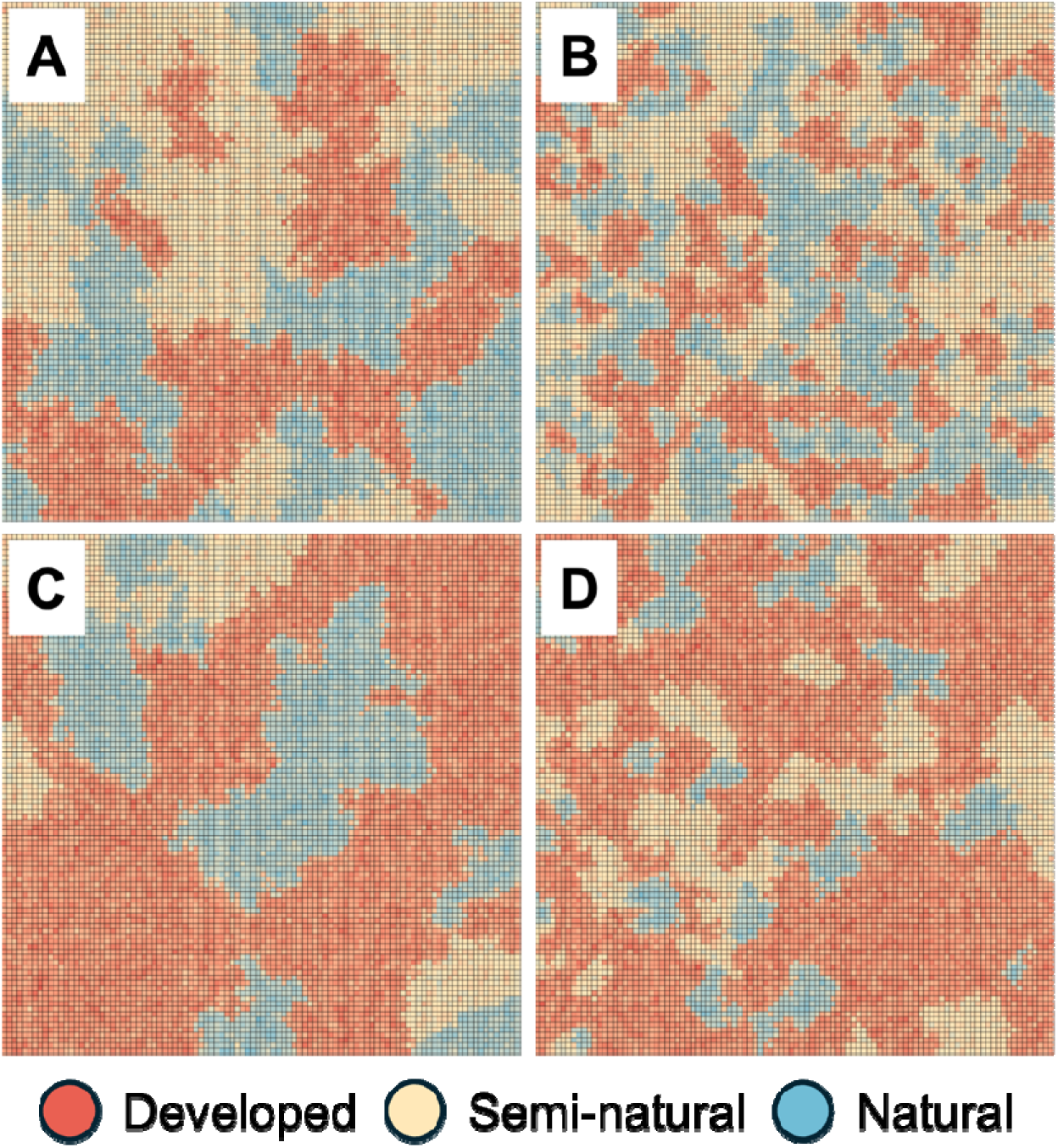
Four example simulated landscapes with different parameterizations. A) A landscape with an even proportion of habitat types with low patchiness. B) A landscape with an even proportion of habitat types with high patchiness. C) A landscape with a higher proportion of developed habitat with low patchiness. D) A landscape with a higher proportion of developed habitat with high patchiness.

### Simulating conservation scenarios

We created two additional functions that took the generated landscapes as input and performed two types of modifications. Both functions converted land from one classification to another, but varied in the locations where land was changed. For this simulation, we changed land types from developed to natural to mimic land acquisition and restoration. The first function converted developed land into natural land along the edges of existing natural land, essentially expanding the edges, and thus made the core habitat larger. This is done by identifying which cells in a developed patch share an edge with a natural patch and then converting a number of these cells based on an input parameter. The other function identified cells in developed patches that were furthest from the edges and then converted a number of these cells based on an input parameter. This scenario was meant to represent the habitat restoration approach of creating stepping stones between natural lands (Saura et al. 2014). Examples of landscape modifications are found in the supplement and the online tool described in the supplementary materials (Fig. S4). The complete set of parameters tested is presented in Table 1.

### Generating additional local and landscape-wide stress

The primary motivation of this simulation is not to explore the role of landscape alone, but also to ask how simulated populations respond to additional stressors when in different landscape types. Additional stressors were classified based on their area of impact, either locally or across the entire landscape. We only added local stressors to semi-natural landscapes, and they acted at the level of the initial patch assignment, where the same negative value was applied to a patch of semi-natural cells. This design resulted in variation in the intensity of local-level stressors across semi-natural cells in one landscape. The magnitude of additional local stress for each patch was randomly drawn from a gamma distribution (multiplied by −1), where values cannot be positive. The rate and shape parameters for the gamma distribution were calculated using moment matching, where the mean of the distribution is a parameter that is varied (Table 1).

Additional landscape-wide stress impacted all cells equally, regardless of land use type, and was intended to represent large-scale “good” and “bad” weather years (such as those generated by ENSO cycles). This parameter was set up as an oscillating sine wave, with a wavelength of six years and an amplitude of 0.5. This effect is of the same magnitude as natural land, and thus, in this simulation, the value of a good weather year is equivalent to that of high-quality habitat. When this parameter was set to the control (of no additional landscape-wide stress), this function oscillated between 0.5 and −0.5 every 6 years. When this condition was set to higher values, a negative trend of varying magnitudes was imposed on this sine function, which caused the oscillating function to worsen over time (following a 20-year burn-in period). A comprehensive description of the local and landscape-wide conditions affecting populations can be found in Table 1.

### Dispersal

After the effects of the environment on population growth were applied to every cell in a time step, the dispersal stage was initiated. For each cell, the percentage of the population emigrating was informed by dispersal parameters (which contribute to a dispersal function). The key parameters in this function were the abundance of the current cell, the habitat quality of the current cell, and two bounding parameters that describe the asymptotes of this function. Lower habitat quality in the current cell or a higher abundance promoted dispersal to other cells (with total dispersal bounded by the asymptotes). Once the percentage of individuals emigrating from a cell is determined, where they go is related to a dispersal distance parameter. When this parameter was set to 1, they could only move within the 3×3 matrix with the current cell as its center, whereas a parameter of 10 allowed them to access cells in a 21×21 area. Closer cells were more likely to be moved to, with the probability being inversely related to distance.

Specifically, we determined the probability that each cell will gain individuals from each of the other cells in the dispersal area (8 potential cells at the lowest setting, 441 at the highest). This was coded as a three-dimensional array, where x and y are the cells in the dispersal neighborhood, and this is repeated z times for all cells in the landscape. For instance, at the lowest setting, x and y are a 9-cell array (with the reference cell in the middle). To determine the number of individuals that the reference cell would gain, we multiplied the emigrating population of each of the eight cells by the inverse distance to the reference cell and added these values together. This final sum became the rate parameter for a single draw from a Poisson distribution. This calculation is performed simultaneously across all 9-cell arrays in the z-dimension. Once the number of individuals each cell gains was determined, the number of individuals a cell loses was calculated and made into a loss matrix. These matrices were then added simultaneously to the population size matrix created after applying environmental effects. The population size after dispersal became the end population size for that year. The functions for dispersal are presented in the supplement (Fig. S5). The complete set of parameters tested can be found in Table 1.

### Output variables

The simulation was run a total of 58,500 times, representing the number of parameter combinations and replications. For each run, we saved the number of times a cell reached zero individuals at any point during the 50-year observation period (extirpated), the number of times an extirpated cell later reached an abundance above the Allee effect threshold (recolonized), and the average abundance in semi-natural cells. We used the extirpation and recolonization metrics to calculate the probability that cells are extirpated and the probability that they are successfully recolonized. We did not count recolonization events in which populations never reached the Allee threshold below which we imposed a negative population growth rate in the simulation (Fig. S1). Our primary metric for inference was calculated using the following equation, which describes the number of cells that are expected to be extirpated (after accounting for recolonization) over 50 years of observing the simulation’s behavior.

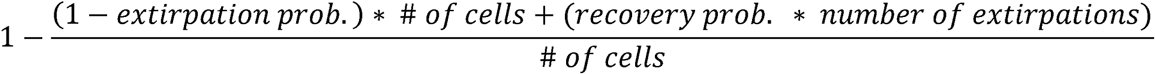

## Results

The number of extirpated cells in a landscape decreased when simulations included more natural habitat and increased with greater development. When the landscape was mainly composed of one of these two habitat types, the number of extirpations essentially matched the number of cells that are either developed or natural (inverse relationship) (Fig. 2A,C). The relationship between extirpations and semi-natural habitat was more complex (Fig. 2B). As the proportion of semi-natural habitat increased between different iterations, the number of populations quickly grew (extirpations decreased) until about one-third of the landscape was semi-natural, after which the effect plateaued and even showed a tendency to reverse with increasing extirpations at higher levels of semi-natural habitat. These effects interacted with dispersal ability.

**Figure 2.**
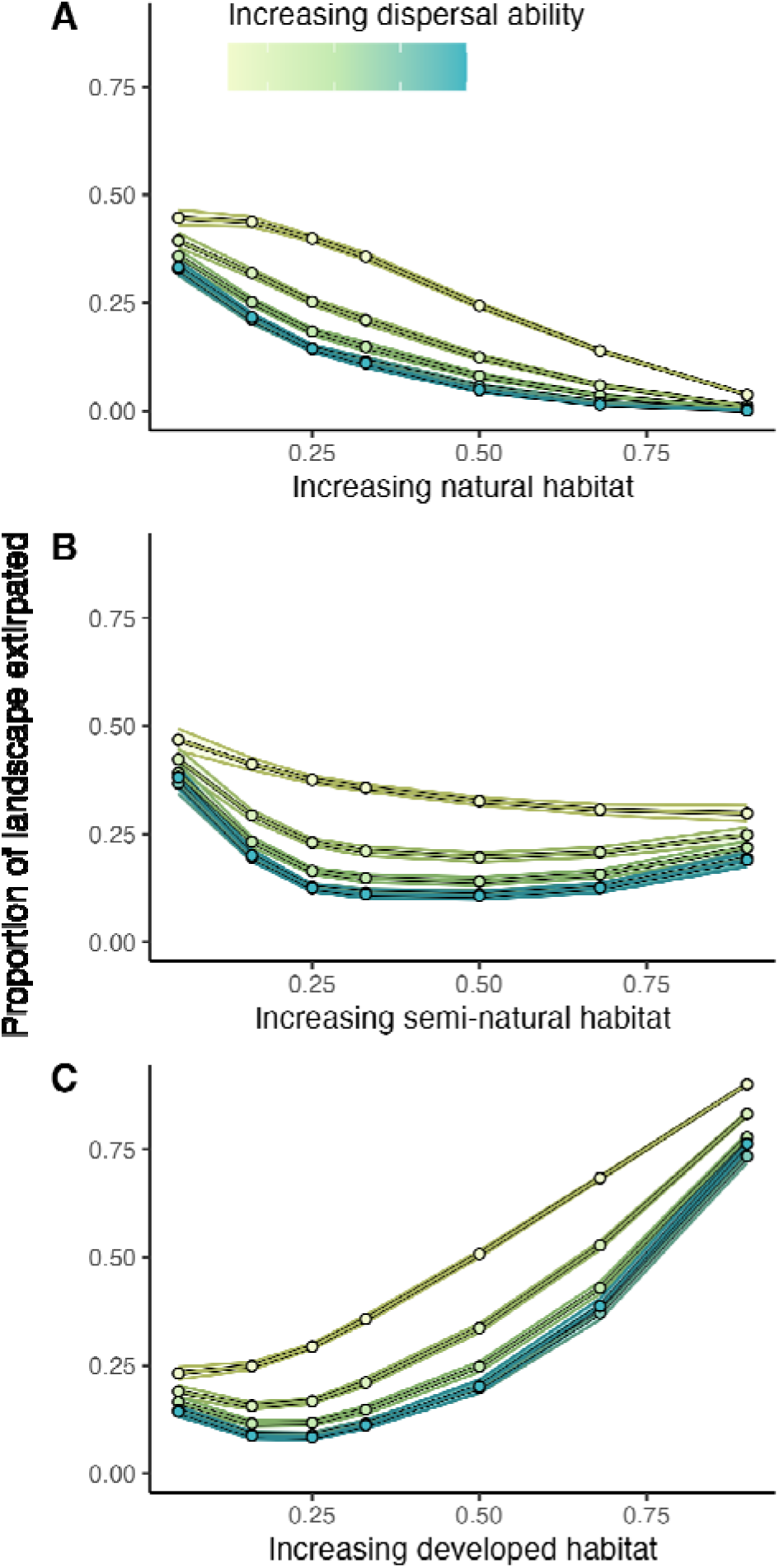
The effects of habitat quality and dispersal ability on the average number of extirpations across the simulations. Panel A shows the impact of increasing natural habitat, Panel B shows increasing semi-natural habitat, and Panel C shows increasing developed habitat. The effects shown are the mean effects after averaging over all other effects. Points and lines represent the mean effect and are colored by dispersal ability. The bands surrounding the lines indicate bootstrapped 95% confidence intervals.

Metapopulations in simulations with more dispersive behavior were more resilient to increased development, occupying about 20% more of the landscape until roughly one-third of it was developed, at which point population density declined rapidly (Fig. 2 A,C). Afterward, the number of populations mostly tracked the amount of usable habitat (natural and semi-natural). This contrasted with simulations of less dispersive dynamics, which closely followed the available habitat (Fig. 2B). Simulations with more semi-natural habitats benefited all species, but the benefit was much greater for dispersive species. For example, when the landscape was 25% semi-natural, approximately 40% of the landscape was extirpated for non-dispersive species, but only about 10% for highly dispersive ones. We also found that species performed between 15 and 25% better in patchier landscapes, and that dispersive species fared better in particular (Fig. 3).

**Figure 3.**
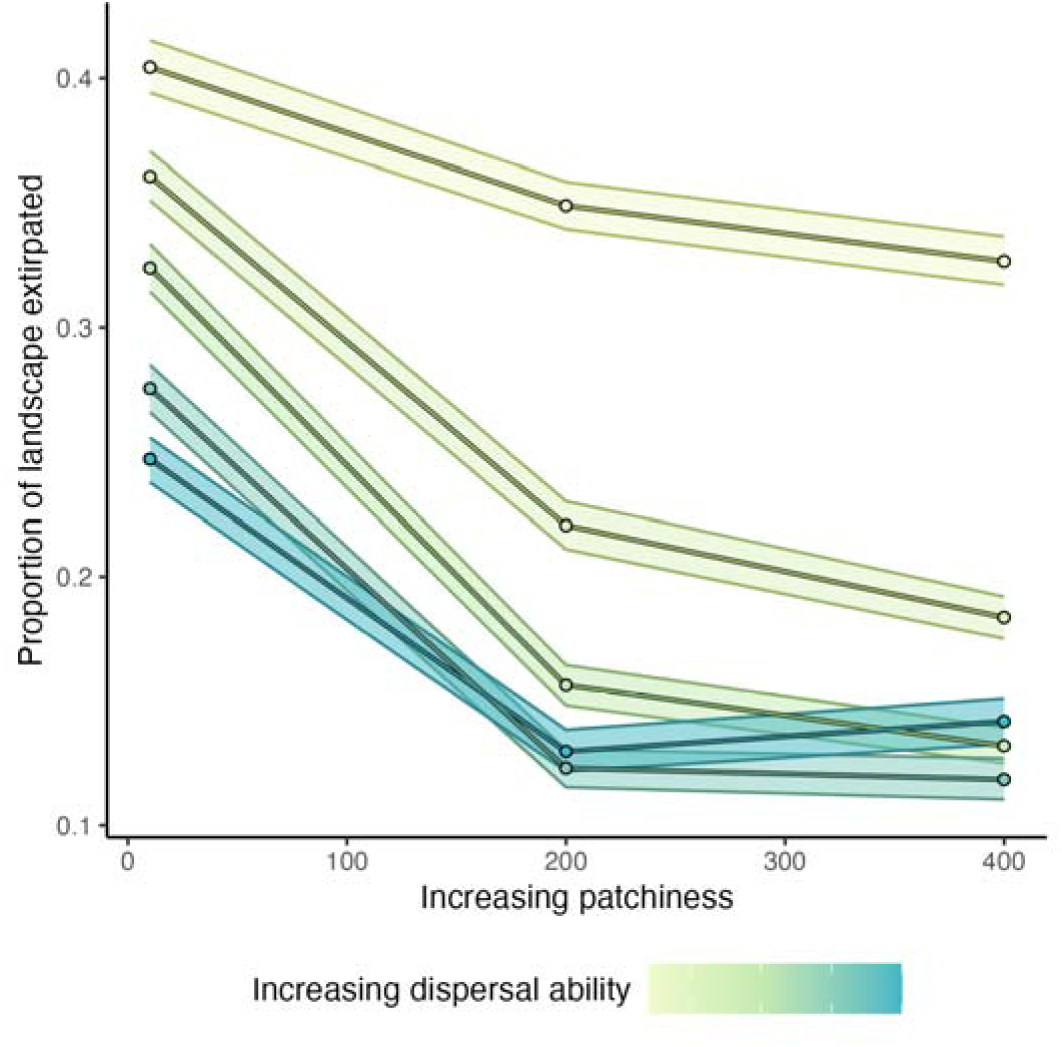
The effects of patchiness and dispersal ability on the average number of extirpations across the simulations. The effects shown are the mean effects after averaging over all other effects. Points and lines represent the mean effect and are colored by dispersal ability. The bands surrounding the lines indicate bootstrapped 95% confidence intervals based on bootstrapping.

The number of extirpations in a landscape also increased when comparing simulations with increasing intensities of additional stressors, which have either a local or landscape-wide effect in addition to the impact of the landscape itself (Fig. 4). The relationship between both types of additional stress and extirpations varied with dispersal. When there was no population stress (other than the landscape itself), the more dispersive species occupied about 20% more of the landscape. As local stress increased, both dispersive and non-dispersive species declined, but the rate of extirpation was higher for non-dispersive species. The change from moderate to high local stress resulted in a 34% increase in extirpations for non-dispersive species, but only a 21% change for very dispersive species (Fig. 4A). The opposite is true in response to regional stress (that impacted all cells), where the rate of extirpation is higher for more dispersive species (Fig. 4B). Specifically, shifts from moderate to high regional stress results in a decline of 10% for non-dispersive species, but a decline of 15% for very dispersive species.

**Figure 4.**
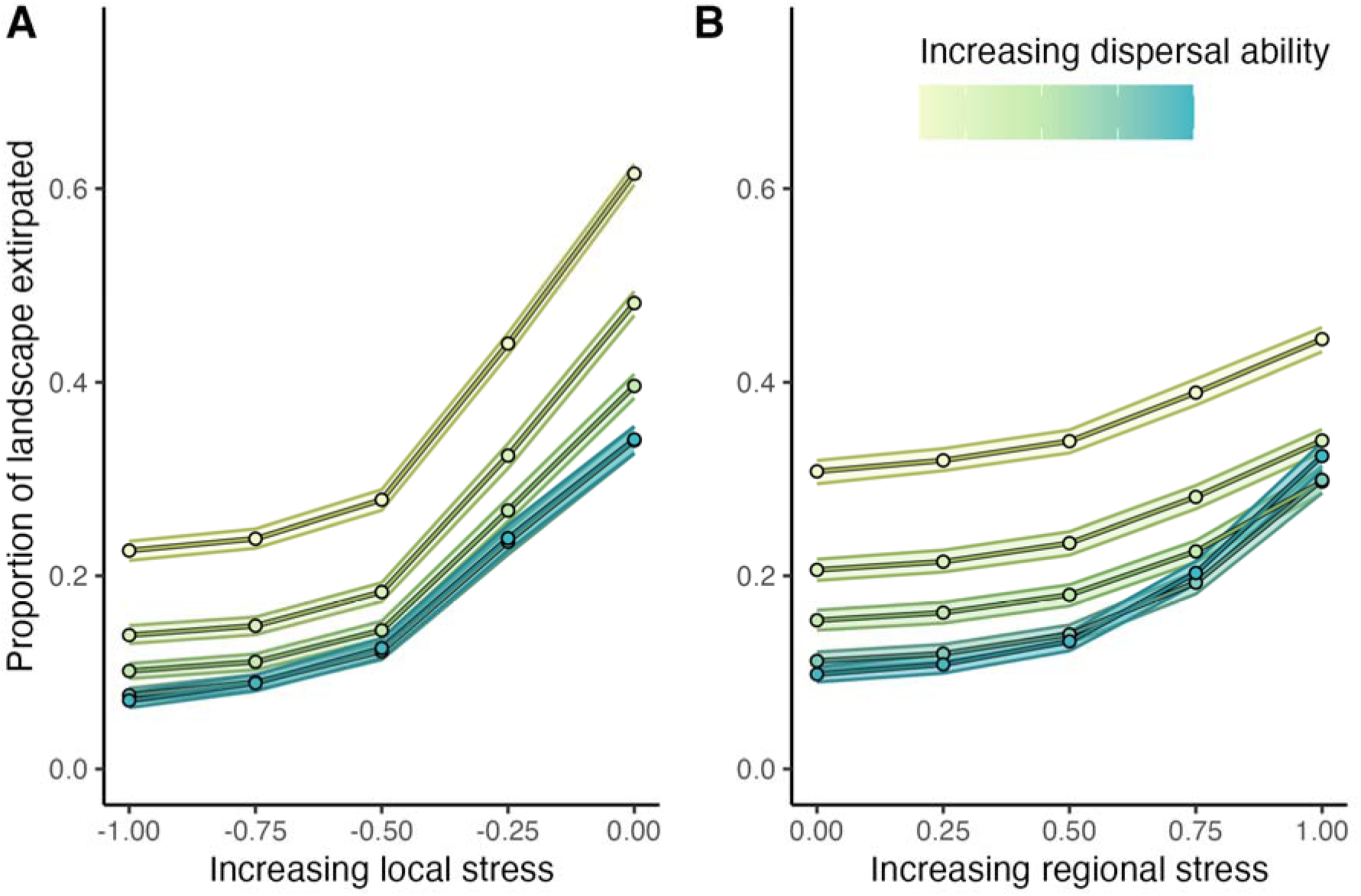
The effects of landscape stressors and dispersal ability on the average number of extirpations across the simulations. Panel A shows the effects of local stressors associated with semi-natural habitat, and Panel B shows stressors that impact the entire landscape. The effects shown are the mean effects after averaging over all other effects. Points and lines represent the mean effect and are colored by dispersal ability. The bands surrounding the lines indicate bootstrapped 95% confidence intervals.

The number of extirpations over the entire simulation also revealed interactive effects between the landscape, combined local and landscape-wide stress, and dispersal capability, shown by the curved contours in Figure 5. When 50% or more of the landscape was comprised of natural habitat, most species, regardless of dispersal ability, were robust to additional stressors (Fig. 5 A-C). The more natural habitat, the better, but all species, even those with little dispersal ability, occupied a large portion of the landscape. When 50% or more of the landscape consisted of semi-natural habitat, more interactions emerged, mainly when the landscape comprised between 50% and 67% semi-natural habitat (Fig. 5 D,E). In these landscapes, a greater ability to disperse resulted in more resilience to moderate amounts of additional landscape stressors. In both instances, simulations with non-dispersive species occupy 20% less of the landscape. As combined stress continued to grow, the dispersive species advantage decreased, and the number of extirpations was more related to the amount of combined stress, and not dispersal (Fig. 5D-F). Landscapes that were primarily developed resulted in the most extreme interactive effects (Fig. 5 G-I). For instance, when the landscape is 67% developed, a non-dispersive species (on the left side of Fig. 5H) declines by 30% when moving from no additional stress to the most extreme case. However, a dispersive species (on the right side of Fig. 5H) declines by 65% when moving from the no additional condition scenario to the extreme additional stress condition. In the most extreme simulation, where 90% of the landscape is developed, there is almost no difference between dispersive and non-dispersive species, under high amounts of additional stress (Fig. 5I).

**Figure 5.**
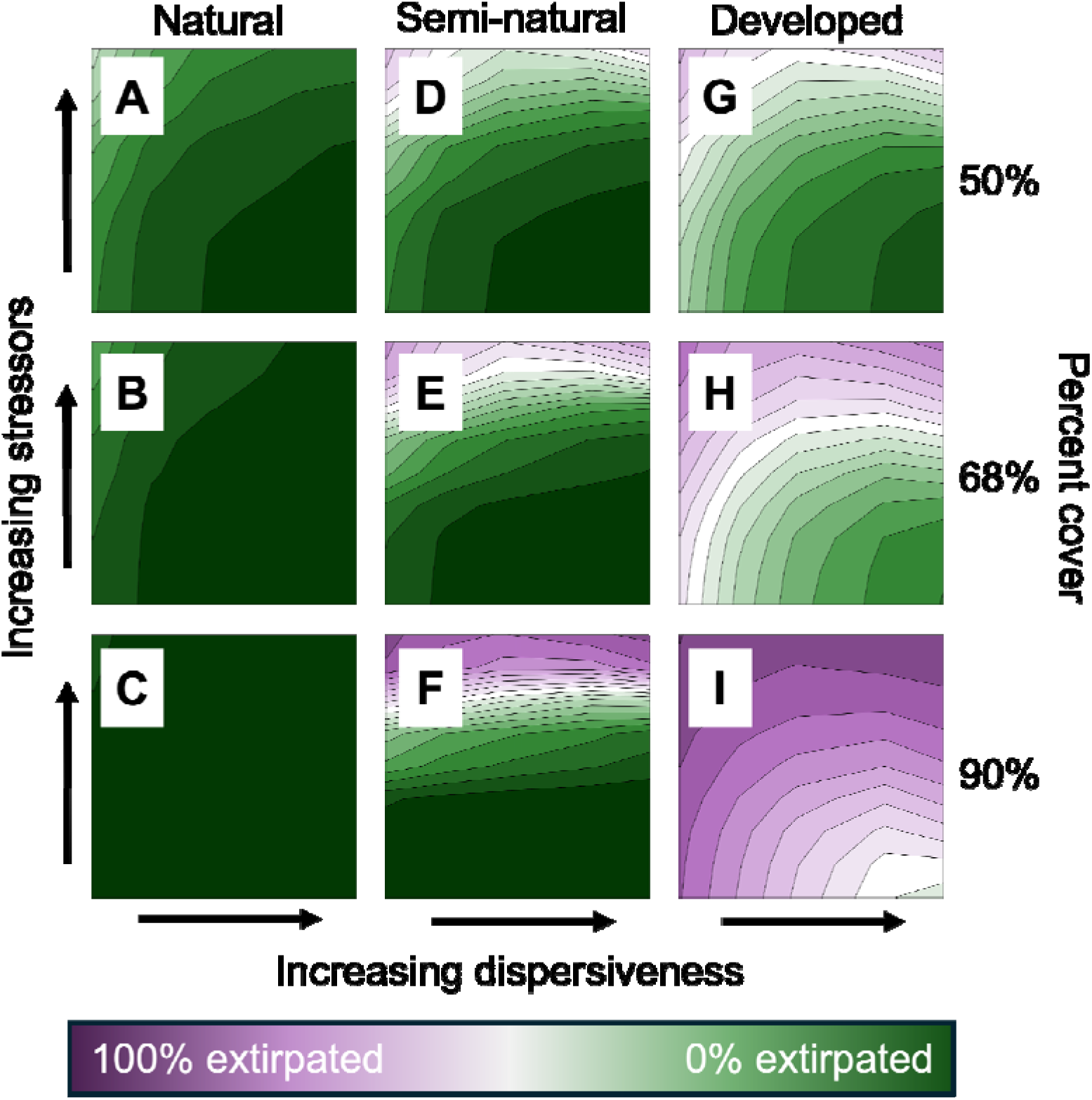
The interactive effects of habitat quality, landscape stress, and dispersal ability on the average number of extirpations across the simulations. Within each panel, the x-axis represents increasing dispersal ability, and the y-axis represents increasing stressors (local and regional combined). Comparison across panels shows the effects of habitat quality. Panels in the left column show when the landscape is primarily natural, panels in the middle show when the landscape is predominantly semi-natural, and panels on the right show when the landscape is primarily developed. As panels move down, the proportion of that cover type increases. Contours and colors represent the average number of extirpations across the simulations, after averaging over the effects not shown. Each contour line represents a change in 5% extirpation.

Interactions between dispersiveness and habitat were not only apparent under simulated stress, but also under simulated conservation. Both simulated conservation scenarios (connectivity versus core habitat) resulted in fewer extirpations; however, increasing connectivity had a non-linear positive effect on dispersive species in non-patchy landscapes. For a mostly non-dispersive species in a non-patchy landscape, the difference between no additional conservation and 30% habitat restoration was an improvement in occupancy of 20%; however, this increase was closer to 40% for highly dispersive species (Fig. 6A). Increasing core habitat was also beneficial; however, the relationship was essentially linear with the amount of new habitat added, regardless of dispersal ability (Fig. 6C). When the simulated landscape was patchier the differences between connectivity and core habitat were not as strong. Both increased approximately linearly with the amount of new habitat being added, with little interaction between dispersiveness (Fig. 6B,D).

**Figure 6.**
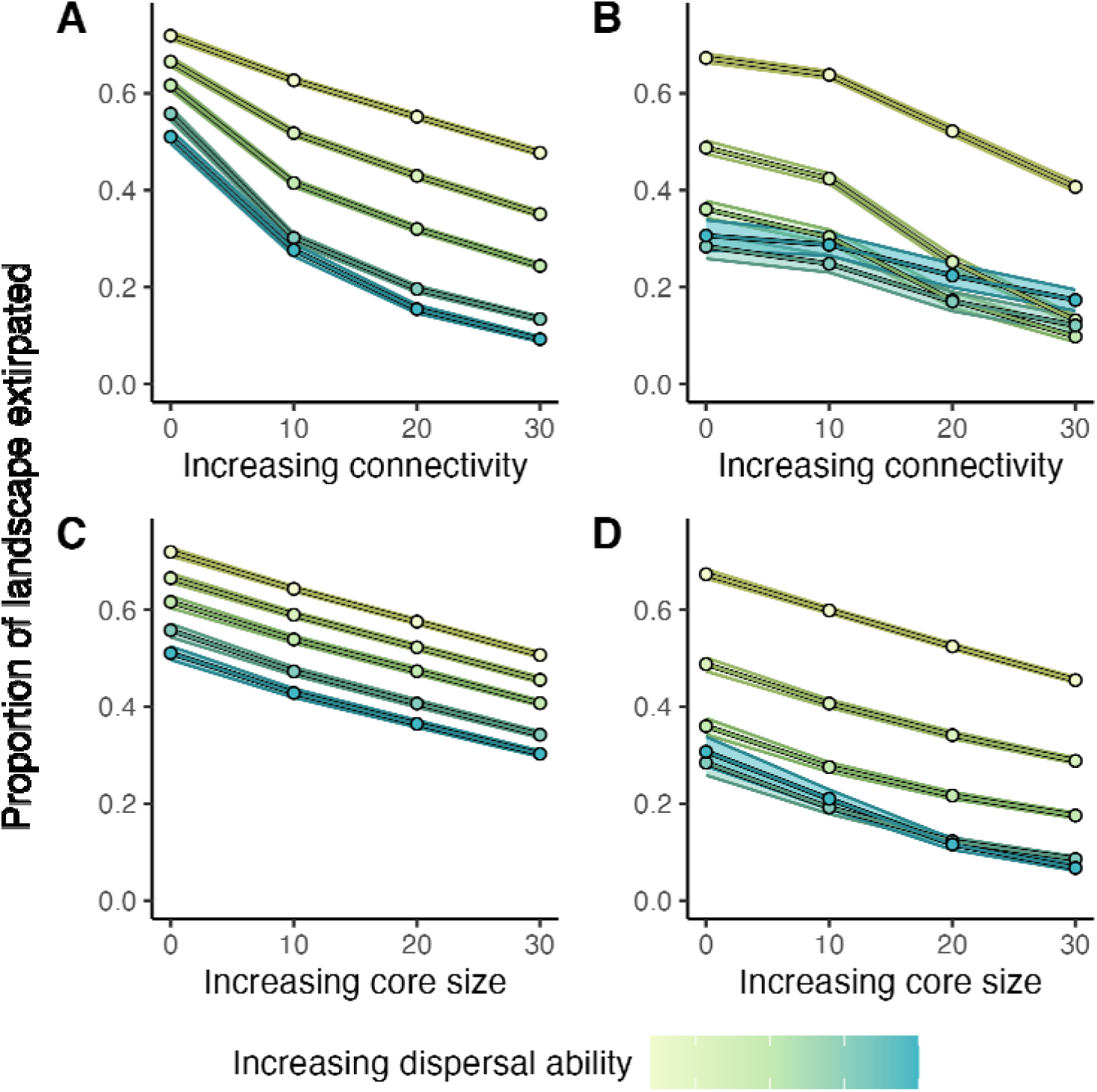
The effects of landscape restoration and dispersal ability on the average number of extirpations across the simulations. Panel A shows the effects of enhancing connectivity between existing patches in a non-patchy landscape. Panel B shows the impact of enhancing connectivity between existing patches in a patchy landscape. Panel C shows the effects of increasing existing natural habitat in a non-patchy landscape. Panel D shows the effects of increasing existing natural habitat in a patchy landscape. The bands surrounding the lines indicate bootstrapped 95% confidence intervals.

In addition to landscape extirpations, we also tracked the average abundance of individuals in semi-natural areas. Overall, this metric interacted with dispersal ability less than extirpations did, and because of this, figures are primarily presented in the supplement (Fig. S6, 7). We found no difference in abundance between dispersal abilities in response to landscape patchiness (Fig. S6). We also found no differences in average abundance between dispersal groups in response to landscape-wide stress (Fig. S7A). We found a slight effect of local stressors, where additional local stress in semi-natural landscapes was more detrimental to non-dispersive species (Fig. S7B). The variable with the most substantial impact was landscape cover, where increases in natural and semi-natural areas disproportionately benefited and increases in development land disproportionately hurt more dispersive species (Fig. 7). This effect was such that, unlike patterns of extirpations, conditions were observed that are clearly worse for dispersive species than non-dispersive ones.

**Figure 7.**
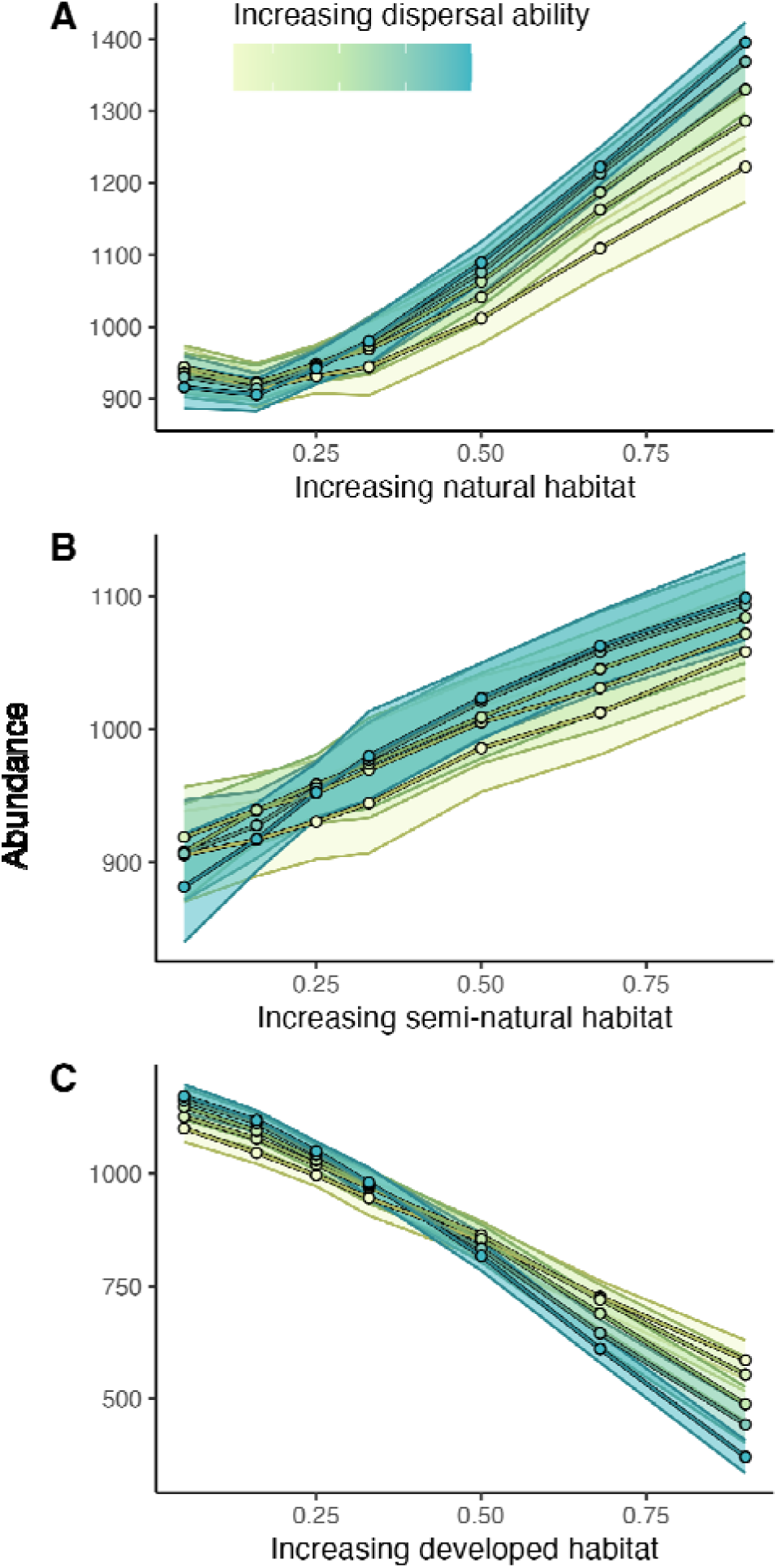
The effects of habitat quality and dispersal ability on the average abundance in semi-natural cells across the simulations. Panel A shows the impact of increasing natural habitat, Panel B shows increasing semi-natural habitat, and Panel C shows increasing developed habitat. The effects shown are the mean effects after averaging over all other effects. Points and lines represent the mean effect and are colored by dispersal ability. The bands surrounding the lines indicate bootstrapped 95% confidence intervals.

## Discussion

Many populations of wild plants and animals are facing concurrent Anthropogenic threats acting at different spatial scales (Brown et al. 2013, Halsch et al. 2025), and we can expect that metapopulation outcomes will depend on the magnitude of stressors and species-specific variation in dispersal ability. Understanding these dependencies may, in part, explain the observed declines of widespread insect species, which are exposed to more landscape stressors than range-restricted species (Van Dyck et al. 2009, Edwards et al. 2025). Here, we developed a spatially explicit metapopulation simulation to simultaneously explore how multiple axes of stress, from landscape composition to local and landscape-wide stressors, and dispersal behavior, impact metapopulation persistence. We found that responses to different categories of stressors were expectedly negative, but that they interacted with dispersal and that expectations for population trajectories may indeed vary with dispersal. These findings have implications for understanding patterns of insect declines, as well as for insect conservation, since strategies to help widespread insects will likely need to consider a larger scale than those for range-limited species.

The primary objective of this study was to gain insight into the nature of the decline of widespread insect species, which include the monarch butterfly and other species in North America that have been reported to have declining populations despite, in many cases, geographic ranges that span multiple broad geographic regions (Forister et al. 2023b, Edwards et al. 2025). We hypothesized that widespread species sample more of the landscape, and because of this, in a heavily modified landscape, they are exposed to more cumulative stress than non-dispersive species. This was not the case for most of the combinations of conditions that we explored. In nearly all combinations of land use compositions and additional stressors, we found that non-dispersive species performed worse, or in extreme cases, about the same as dispersive ones. This does not mean that dispersive species are not at risk, as long-term data indicate that many are in decline (Van Dyck et al. 2009, Forister et al. 2021); instead, it invites consideration of the areas in our simulated world that most closely reflect the patterns of decline of both non-dispersive and dispersive species. The conditions where this was seen were in landscapes that were highly developed, where most of the landscape provides no resources for synthetic populations. It is in these landscapes where dispersive species are reduced to similar occupancy as non-dispersive ones. This is likely due to the reduction of connectivity. Connections between source populations reduce distances to other high-quality patches and can stabilize the network through the effects of long-range dispersal (Howe et al. 1991, Johst et al. 2002, Bowler and Benton 2009). In primarily developed landscapes, the semi-natural spaces become fewer and more critical, and as additional pressures increase in these spaces, this is especially detrimental to dispersive species.

This result may provide some insight into the contemporary loss of butterflies in highly converted landscapes (Van Dyck et al. 2009, Forister et al. 2010). We consider the case of butterflies in the Central Valley of California as an example of such a landscape; however, the results apply to other, similarly modified landscapes. The Central Valley is a highly converted landscape, where the legacy of centuries of land use change is now being met with additional factors like invasive species, disease, pesticides, and climate change (Forister et al. 2010, MacLean et al. 2018, Halsch et al. 2020). This is precisely the type of landscape where, based on our results, we should expect to see rates of decline that are comparable regardless of dispersal range. It may be the case that the historical loss of habitat over centuries had greater impacts on non-dispersive species (as predicted by the simulation), and that contemporary dispersive insects are only in significant decline now in response to additional stressors like climate change, introduced species (and pathogens), and pesticide exposure. In other words, the decline of many range-restricted species may have occurred before their geographic distributions and habitat preferences were completely understood (Fattorini 2011, Habel et al. 2016, Forister et al. 2023a).

While the extreme landscapes yield extreme results for dispersive species, it remains true that this simulation demonstrates that being dispersive is mostly beneficial, even if it means being exposed to the additional threats of a semi-natural landscape. One likely contributing factor to this finding is the landscape generation process itself. The initial patches from which each landscape was built were placed randomly, which, on average, results in well-connected and stable landscapes (Grilli et al. 2015). For instance, even if a landscape is 2/3 developed, the remaining 1/3 will be more evenly distributed across the matrix, resulting in a landscape that may benefit dispersive species (Howe et al. 1991). The distribution of natural land in human-modified landscapes is not random, and it is possible that specific landscapes can be constructed to hurt dispersive species even more than non-dispersive ones. Such a study would be an excellent follow-up to these results, especially if it were designed based on the landscape compositions of real focal landscapes. Still, this simulation demonstrates that under many conditions, a general expectation is that dispersive species are more stable as a metapopulation network than non-dispersive species.

Another explanation for our results differing from our expectations may result from the primary metric of inference: the percent of the landscape extirpated. This metric summarized the entire landscape and, based on extirpation and recolonization rates, calculated the expected number of cells expected to be occupied at equilibrium. This metric is thus all-knowing and incorporates information from the entire landscape to assess extinction. This is not the same type of information that is provided by long-term monitoring programs, which would be more reflected by the abundance of a single cell in the simulation. Furthermore, many large-scale monitoring programs are biased in their location, as they often oversample areas near development (Dunn et al. 2005, Geldmann et al. 2016). For this reason, we also tracked the average abundance of species in semi-natural cells. We found that there are indeed cases where dispersive species decline at the same rate or at greater rates than non-dispersive species. The factor that generated the most significant declines of dispersive species (relative to non-dispersive ones) was the development of land. This is likely a further demonstration of the previously noted importance of connectivity for dispersive species (Johst et al. 2002) and that the transition to a landscape entirely unusable has detrimental effects on dispersive species.

This result also demonstrates that while monitoring data may detect greater reductions in the abundance of widespread species, this does not necessarily reflect a greater risk to the entire metapopulation. This suggests that even in cases where dispersive species experience greater losses of individuals across the whole meta-population, they are still at less total risk of complete extirpation. While this may be the base expectation, this finding should be taken with caution.

Firstly, many charismatic widespread insects, such as the monarch butterfly (*Danaus plexippus*), are not only highly dispersive but also migratory (Reppert and Roode 2018), and these dynamics are not captured in this simulation. For instance, we did not examine situations where conditions in a subset of the landscape, such as overwintering grounds, exhibit a disproportionate influence on the entire population. In such cases, a model would need to be built to capture this dynamic explicitly, and it may be the case that the impacts on population growth during this one critical phase of their migration are as crucial as the rest of the landscape collectively (Taylor and Hall 2011). Secondly, while under many conditions, non-dispersive species may be at a higher risk of metapopulation extirpation, there are conditions where dispersiveness ceases to matter, which occurs in landscapes that are heavily modified and under additional stressors. Given the landscape composition of many lowland areas and the global threat of climate change, it is not unreasonable to expect that many species in such places are being pushed to their limits and that declines of range-restricted and widespread species should be expected.

While this simulation is informative for understanding observed patterns of insect decline, it can also inform the efficacy of potential interventions. Semi-natural landscapes appear to offer considerable value for bolstering metapopulations, especially for more dispersive species (Howe et al. 1991). While the expansion of high-quality source habitats is beneficial for all species, increasing the coverage of semi-natural habitats can have comparable effects for dispersive species (to a certain extent). In the context of insects, such areas may include pollinator strips and hedgerows in agricultural areas, as well as pollinator gardens in cities, which can provide resources in otherwise substandard areas (Buhk et al. 2018, von Königslöw et al. 2022, Donkersley et al. 2023). This finding is also supported by experimental work, which has shown that open corridors that do not support populations themselves are still capable of facilitating movement between high-quality patches (Haddad and Tewksbury 2005). Our additional habitat restoration iterations also found that building connections between existing high-quality patches can have a disproportionately positive effect on dispersive species. The effect was especially strong in landscapes that are not otherwise patchy, with few existing connections between source populations. Such “stepping stone” habitats are important for long dispersal events, although such habitats need to be of adequate size or quality to be useful (Saura et al. 2014). Together, these results underscore the importance of strategically expanding and connecting semi-natural and high-quality habitats, which can amplify the persistence of dispersive species.

One of the defining features of the Anthropocene is the interconnectedness of threats imposed on natural populations (Breitburg et al. 1998, Pirotta et al. 2022, Halsch et al. 2025). Many landscapes face multiple at once, which is likely crucial for understanding the decline of both local and widespread insects (Yang et al. 2021, Pirotta et al. 2022). Our findings provide a metapopulation perspective on this process and offer further insights into the decline of widespread species. All else being equal, being highly dispersive creates larger and more stable metapopulations. Still, in highly developed landscapes with a legacy of habitat loss, additional stressors like climate change can lead to rapid declines. These results highlight the importance of marginal habitats and how establishing connections can support mobile species (Samways 2007). Many range-limited species may have already been lost before baseline data could be established through monitoring programs. However, many widespread species, while declining, can be supported through thoughtful land conservation and management.

## Acknowledgments

C.A.H. acknowledges the USDA NIFA predoctoral fellowship (2022-67011-36563) for funding this idea and the initial development. M.L.F. thanks the National Science Foundation (DEB-2114793). E.M.G. thanks the National Science Foundation (DEB-2225092).

