## Supplement for "Dispersal mediates metapopulation response to local and regional stressors"

**Dispersal ability mediates responses to local and regional stressors to shape metapopulation persistence**

This file includes:

Description of online resources that demonstrate landscape generation and running the simulation.

Method for power analysis to determine sampling effort

Figure S1. Functions that determine population growth each year.

Figure S2. Demonstration of the landscape modification functions.

Figure S3. Functions that determine dispersal.

Figure S4. The reduction in error in the population estimate as the number of iterations for each parameter combination increases.

Figure S5. Distribution of the ratio of within landscape variance to between-landscape variance when comparing landscapes of the same proportion of land types.

Figure S6. The effects of patchiness and dispersal ability on the average abundance in semi-natural habitats across the simulations.

Figure S7. The effects of landscape stressors and dispersal ability on the average abundance in semi-natural cells across the simulations.

Table S1. The effects of different parameters in the simulation on variability in the simulated answer.

**Description of online resources that demonstrate landscape generation and running the simulation.**

We have built a website to support this project that describes the simulation process, provides a page for generating landscapes with different input parameters, and a page to view the simulation given a subset of parameters. The landscape generator does so in real time and will produce different results given the same input parameters. The simulation running app is pre-rendered to save computational time (repeating it will produce the same results) and only shows a subset of parameters. Both can be found here: https://www.chrishalsch.com/butterfly_landscapes/

**Method for power analysis to determine sampling effort:**

We performed an initial analysis to determine the optimal number of iterations for parameter combinations in the full simulation. This was done by running a reduced version of the simulation where we generated landscapes that were either the most extreme proportions for all three land types (90% developed, semi-natural, or natural) or the most even (where the landscape is 33% of each type). We also ran the extreme and moderate versions for each parameter of interest (dispersal, regional trend, local trend). This reduced simulation was run for each parameter combination and on each landscape ten times and this process was repeated for ten landscapes with the same proportions. This design allowed us to determine the parameters that produce the most variation in results and identify whether within-landscape or between-landscape variation produces more uncertainty. We assessed how well a particular run performed by comparing the result of each single run to the mean across all runs with the same parameters in the same simulated landscape. Essentially, the “true” answer is the value determined when you use the most information from the simulation as the best answer.

We assessed which parameters contributed the most to variability in results using visual and statistical approaches. We performed a rarefaction-style analysis, taking subsets of the ten runs (for every parameter combination) and asking what the average gap is to the “true” answer when only running 2,3, etc., simulations. Doing this, we found that the most frequent increase in reliability in the final answer was running a specific parameter combination more than 3 times (Fig. S5). After this, there is always improvement, but the relationship between the amount of improvement and sampling became linear rather than exponential. To identify which parameters in the simulation contribute the most to variation, we modeled differences from the “true” mean using each parameter in the model (we treated each as a categorical variable). This was done using the glm function in R. We found that the amount of semi-naturalness and dispersal distance had the highest impact on variability. Thus, we elected to focus on these for increased sampling in the full simulation (Table S1). Finally, we compared the within-landscape variance to the between-landscape variance using an anova and found that between-landscape variation is much higher than within-landscape variation (Fig. S6).

Using this information, we elected for the final design where most parameter combinations were run on five randomly generated landscapes (with no repetition within a landscape). Since variability in the final result was most related to the amount of semi-natural habitat, we elected to scale the number of iterations with this variable in the following way:

Semi-natural < 0.1 = 5 iterations; Semi-natural < 0.33 = 8 iterations, Semi naturalness < 0.5 = 12 iterations; Semi naturalness > 0.5 = 15 iterations.

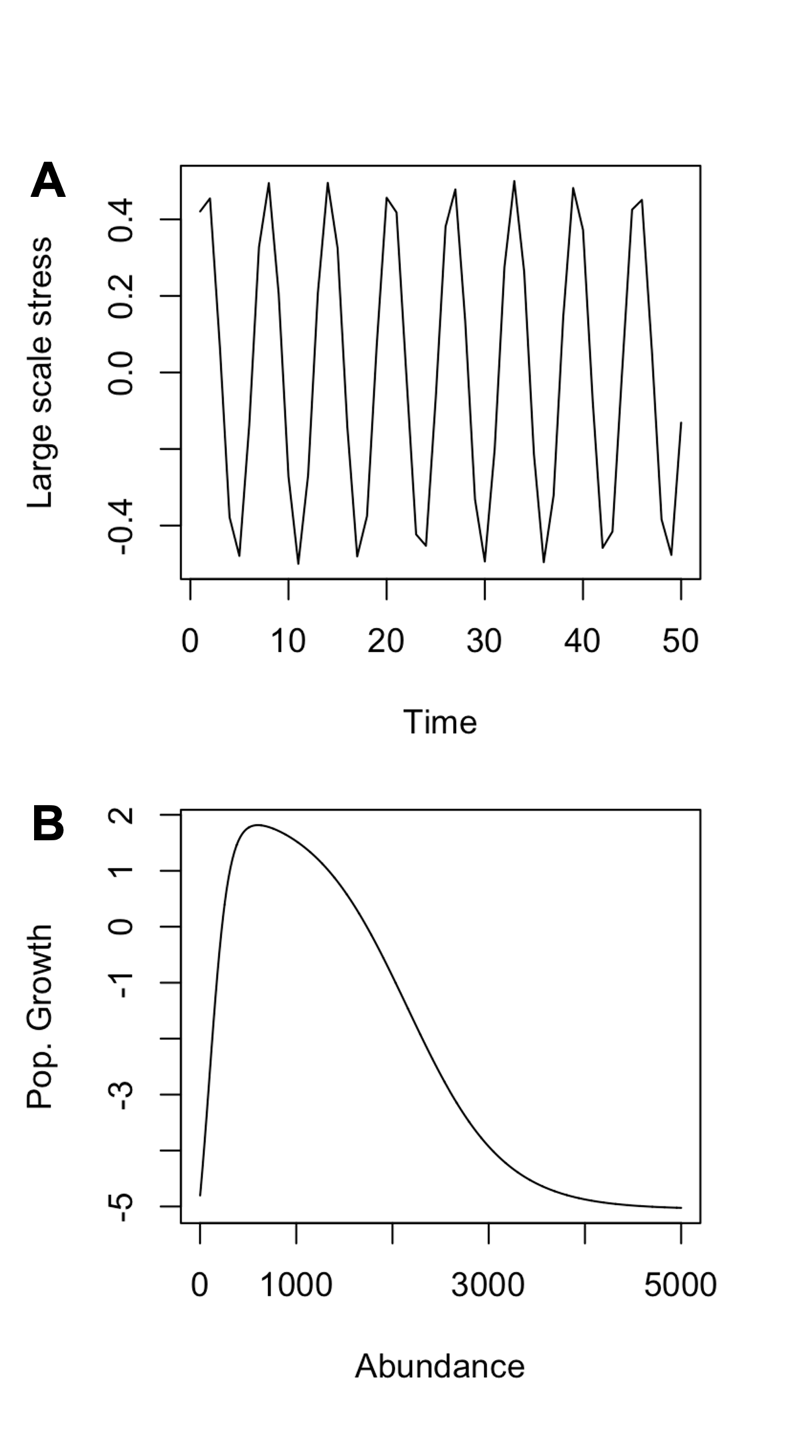

Figure S1. Functions that determine population growth each year. A) Weather fluctuations over the simulation's length (with no trend imposed). Weather impacts all cells equally. B) The function for density dependence in a given cell. Extreme low and high numbers of individuals result in negative population growth.

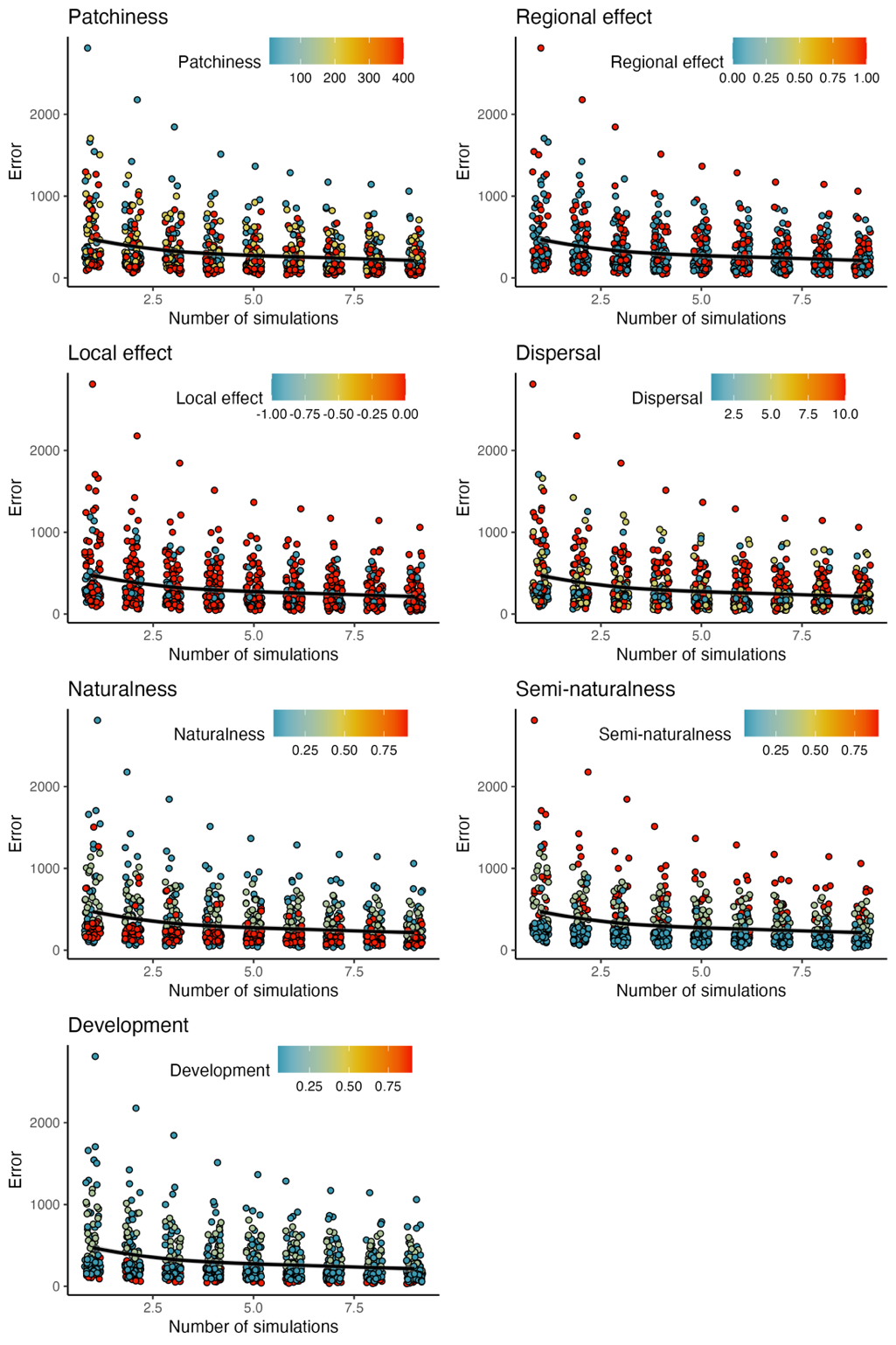

Figure S2. The reduction in error in the population estimate as the number of iterations for each parameter combination increases. Each plot shows the same relationship, but the points are colored by the different tested variables. Points are slightly jittered horizontally to better display data.

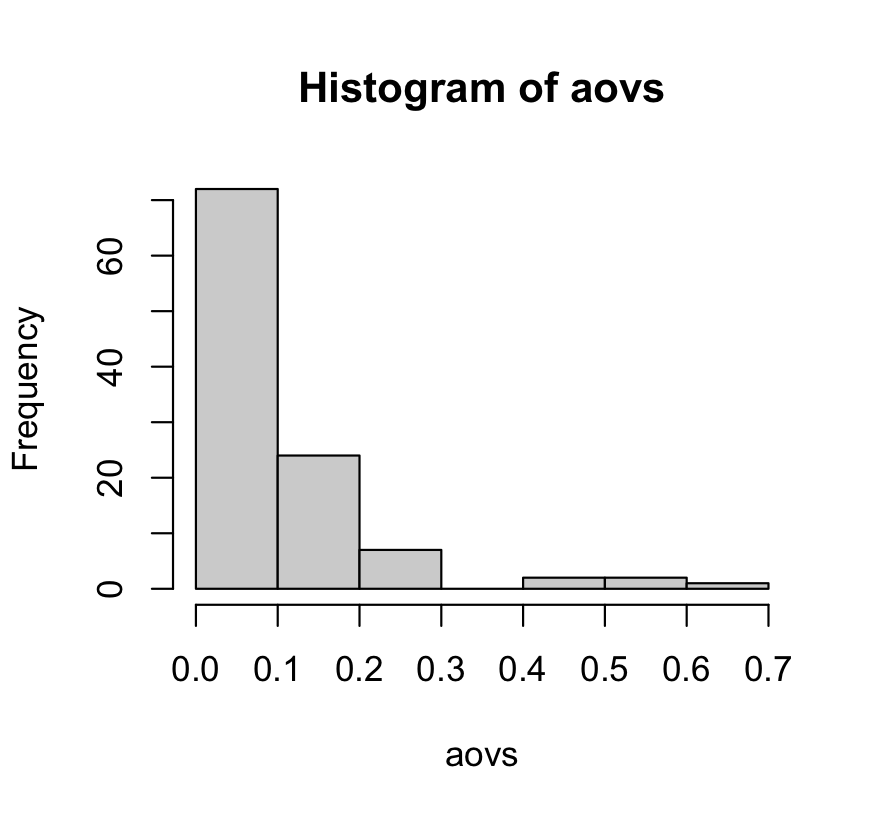

Figure S3. Distribution of the ratio of within landscape variance to between landscape variance on when comparing landscapes of the same proportion of land types. The majority of variation is introduced between landscapes rather than from multiple runs on the same landscape.

.

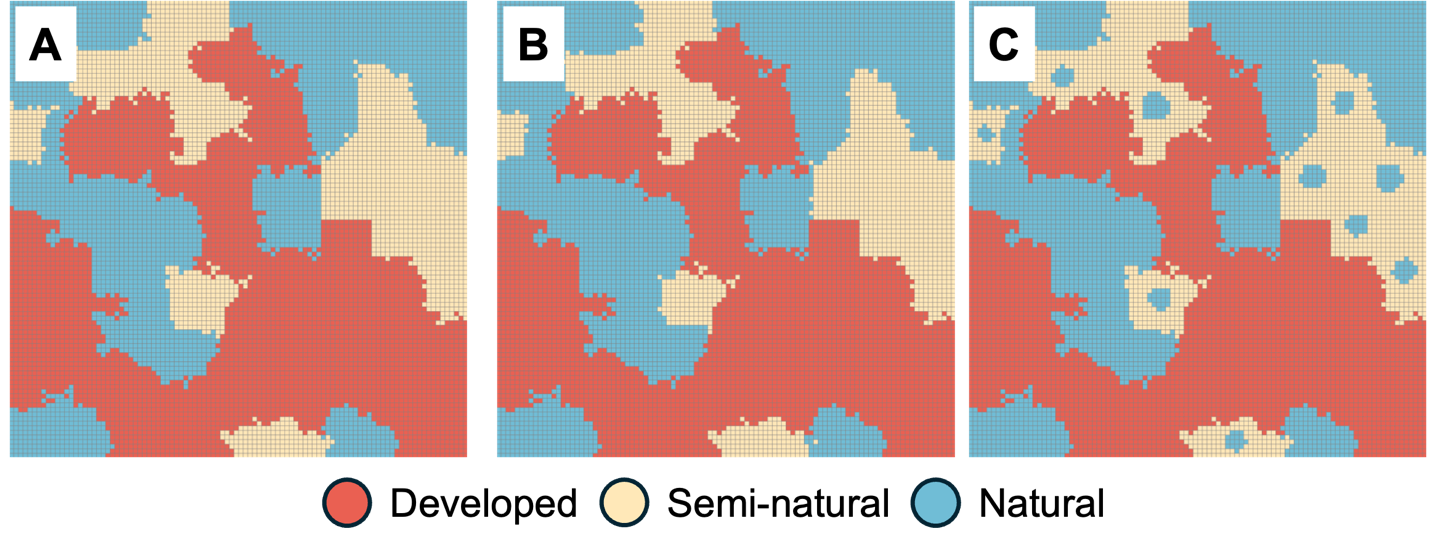

Figure S4. Demonstration of the landscape modification functions. A) The original landscape before any modification. B) The function that turns borders between natural and semi-natural habitats into a natural habitat. C) The function that turns areas in the middle of semi-natural patches into natural habitat. In figures B and C the same number of cells were converted.

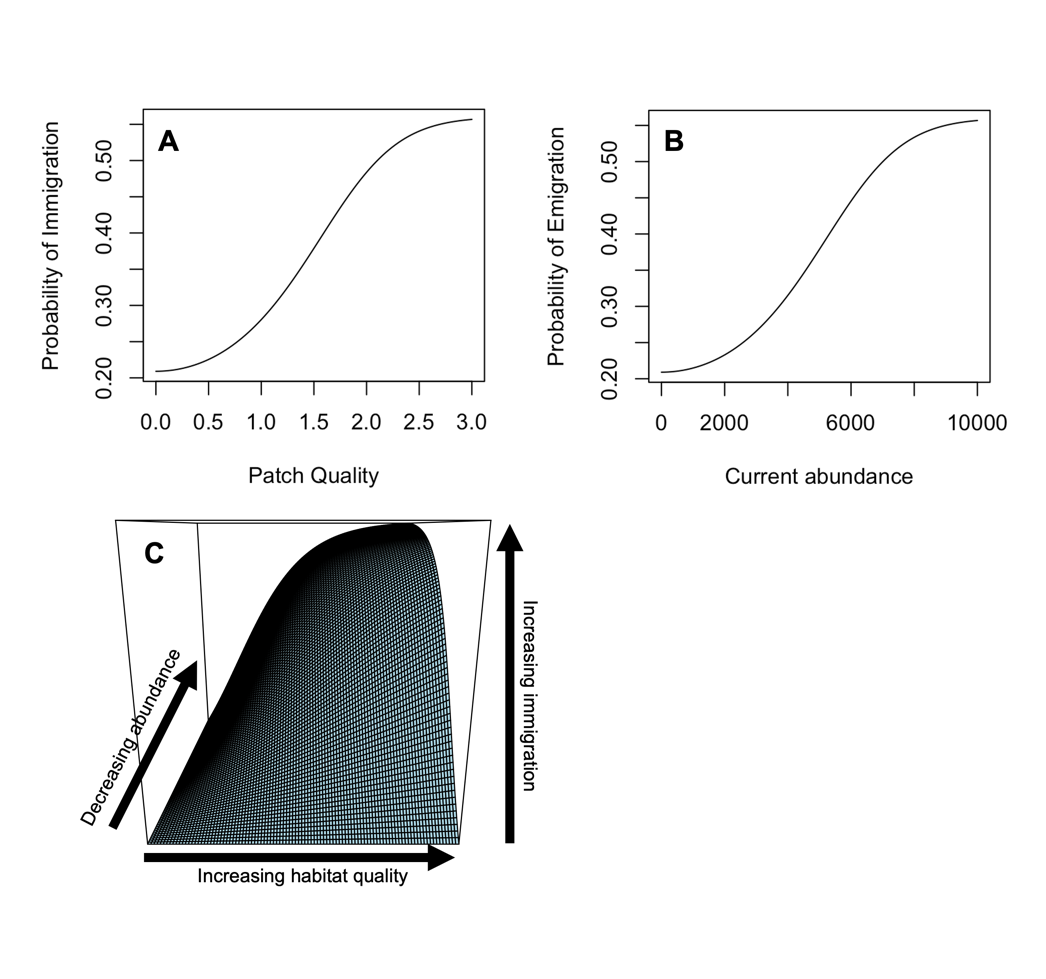

Figure S5. Functions that determine dispersal. A) Individuals favor moving into patches that have a higher patch quality. B) Individuals favor moving from patches with higher abundance. C) The combinations of panels A and B in a single figure.

| 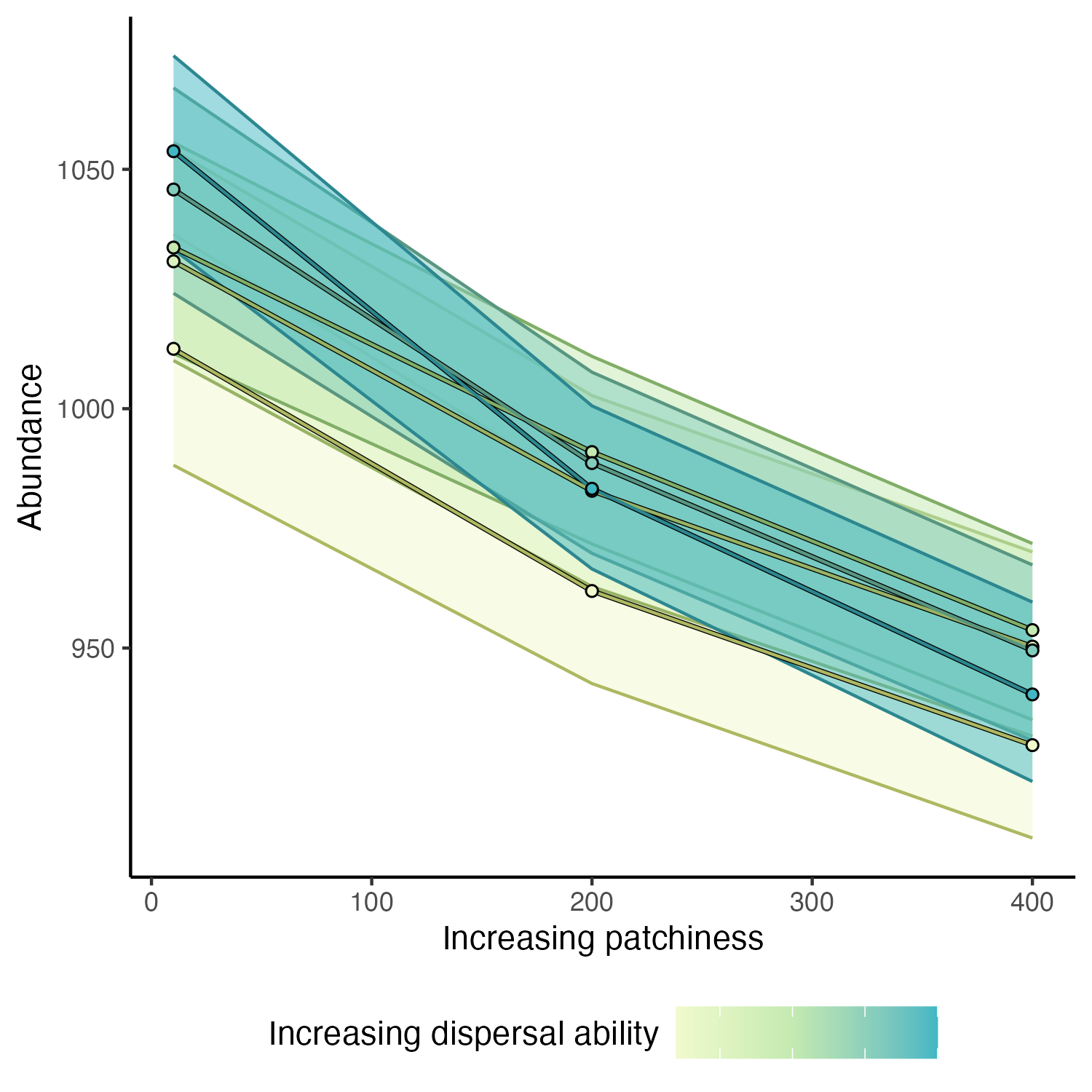  Figure S6. The effects of patchiness and dispersal ability on the average abundance in semi-natural habitats across the simulations. The effects shown are the mean effects after averaging over all other effects. Points and lines represent the mean effect and are colored by dispersal ability. The bands surrounding the lines indicate bootstrapped 95% confidence intervals based on bootstrapping.  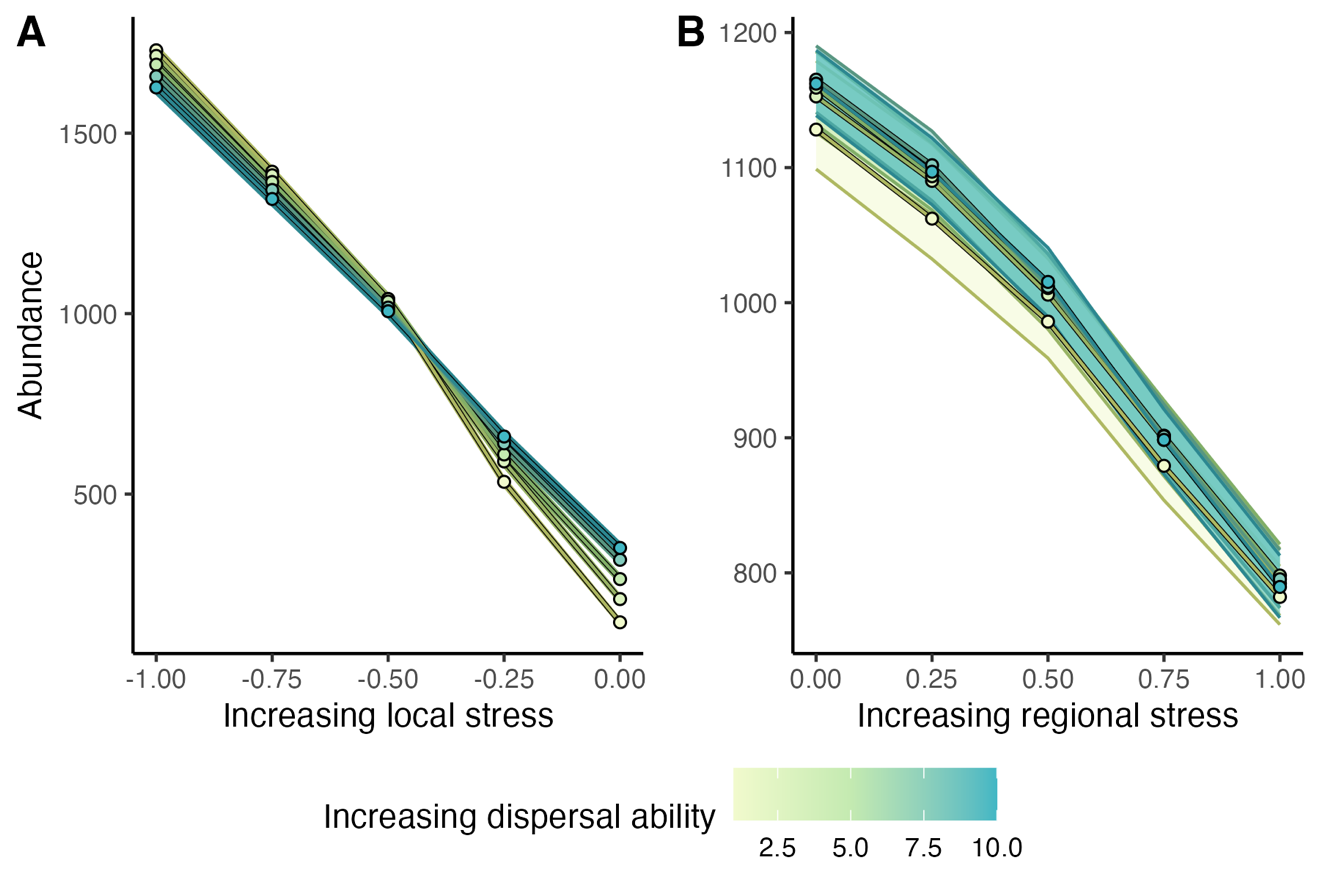  Figure S7. The effects of landscape stressors and dispersal ability on the average abundance in semi-natural cells across the simulations. Panel A shows the effects of local stressors associated with semi-natural habitat, and Panel B shows stressors that impact the entire landscape. The effects shown are the mean effects after averaging over all other effects. Points and lines represent the mean effect and are colored by dispersal ability. The bands surrounding the lines indicate bootstrapped 95% confidence intervals.  Table S1. The effects of different parameters in the simulation on variability in the simulated answer. | | | |
| --- | --- | --- | --- |
| Covariate | | Change in variation | p-value |
| Intercept |  | -60.33 (± 14.09) | >> 0.01 |
| Landscape proportions | 33 % Semi natural | 168.20 (± 10.74) | >> 0.01 |
|  | 90% Semi natural | 216.96 (± 10.74) | >> 0.01 |
|  | 90% Natural | 33.07 (± 10.74) | >> 0.01 |
| Landscape patchiness | Medium patchiness | -41.52 (± 9.31) | >> 0.01 |
|  | Max patchiness | -91.95 (± 9.31) | >> 0.01 |
| Local effect | Max effect | 38.67 (± 9.31) | >> 0.01 |
| Regional effect | Max effect | 26.11 (± 9.31) | >> 0.01 |
| Dispersal distance | Medium dispersal | 32.08 (± 9.31) | >> 0.01 |
|  | Max dispersal | 96.97 (± 9.31) | >> 0.01 |
